# Altered function of the glutamate aspartate transporter GLAST in glioblastoma

**DOI:** 10.1101/287623

**Authors:** Cristina Corbetta, Natalia Di Ianni, Maria Grazia Bruzzone, Monica Patanè, Bianca Pollo, Gabriele Cantini, Manuela Cominelli, Ileana Zucca, Federica Pisati, Pietro Luigi Poliani, Gaetano Finocchiaro, Serena Pellegatta

## Abstract

In glioma patients, high levels of glutamate can cause brain edema and seizures. GLAST, a glutamate-aspartate transporter expressed by astrocytes with a role in glutamate uptake, is highly expressed on the plasma membrane of glioblastoma (GBM) cells, and its expression significantly correlates with shortened patient survival. Here, it was demonstrated that inhibition of GLAST expression limited the progression and invasion of GBM xenografts. Magnetic resonance spectroscopy was used to measure glutamate in GLAST-expressing gliomas showing that these tumors exhibit increased glutamate concentration. Despite their GLAST expression, GBM stem-like cells (GSCs) released rather than took up glutamate due to their lack of Na+/K+-ATPase. Overexpression of Na+/K+-ATPase in these cells restored glutamate uptake and induced apoptosis. The therapeutic relevance of targeting GLAST in gliomas was assessed using the inhibitor UCPH-101. In glioma-bearing mice, a single intratumoral injection of UCPH-101 significantly increased survival by decreasing GLAST expression and inducing apoptosis. Thus, GLAST has a novel role in GBM that appears to have crucial relevance in glutamate trafficking and may thus be a new therapeutic target.

## Introduction

Glutamate is a major cause of brain toxicity associated with glioblastoma (GBM) growth and may be responsible for brain edema as well as other tumor-related symptoms, including seizures. A number of studies have also suggested that glutamate has a crucial role in supporting the invasion and long-distance diffusion of GBM cells (Lyons *et al*, 2007; Ye & Sontheimer, 1999a). These findings indicate that glutamate metabolism and trafficking are profoundly altered in GBM tumors and surrounding normal tissues.

The Na-dependent transporters GLAST (also, EAAT1 or SLC1A3) and GLT-1 are expressed by astrocytes and have a role in glutamate uptake. These transporters help maintain intracellular glutamate levels and prevent glutamate-mediated excitotoxicity in the CNS, and their activity depends on the Na+ electrochemical gradient generated by Na+/K+-ATPase (Attwell *et al*, 1993; Szatkowski *et al*, 1990). Functional reversal of the GLAST transporter in astrocytes has been reported under pathological conditions including ischemia and is responsible for the majority of glutamate release. The collapse of the Na+ gradient that drives glutamate uptake increases the intracellular glutamate concentration, which causes the transporters to reverse their function and thereby release glutamate (Attwell *et al*, 1993; Szatkowski *et al*, 1990; Camacho & Massieu, 2006). Alterations in glutamate transport have been found in glioblastoma cells. For example, these cells cannot take up glutamate due to the absence of GLT-1, which is repressed by the oncogene AEG-1 (Lee *et al*, 2011) or as a result of GLAST mislocalization into the nucleus (Ye *et al*, 1999). Thus far, cystine-glutamate antiporter xCT has been considered the only system through which GBM cells release excitotoxic concentrations of glutamate (Buckingham *et al*, 2011), and this exchanger has also been related to the outcome of patients with GBM (Robert *et al*, 2015). Branched-chain amino acid transaminase 1 (BCAT1) was also proposed to contribute to glutamate production through BCAA catabolism in IDH^wt^ gliomas (Tönjes *et al*, 2013).

Using a culture system that allows the generation and expansion of GBM neurospheres (also defined as glioma stem cells, or GSCs) (Finocchiaro & Pellegatta, 2016), we discovered that, despite their expression of GLAST, GSCs cannot take up glutamate. The down-regulated or absent expression of Na+/K+-ATPase in GBM cells can trigger the altered glutamate transport leading to a decreased glutamate uptake and subsequent extracellular accumulation. Direct correlations between GLAST expression and decreased survival of GBM patients support the clinical relevance of this observation. We also examined the effects of UCPH-101 (Abrahamsen *et al*, 2013; Jensen & Bräuner-Osborne, 2004; Jensen *et al*, 2009), an EAAT1/GLAST blocker that has previously been tested in normal cells (Takaki *et al*, 2012; Tse *et al*, 2014), on glioma cells. GLAST inhibition significantly increased apoptosis in GSCs, but not astrocytes. In vivo, a single intracranial administration of UCPH-101 induced a partial effect. Altogether, these findings provide a foundation supporting translational research aimed at identifying novel GLAST inhibitors that can cross the blood-brain barrier (BBB).

## Results

### GLAST is highly expressed in glioblastoma specimens

Our first evidence that glioma stem-like cells (GSCs) can express GLAST came from gene expression profiling of murine GL261 cells growing as neurospheres (NS) or adherent cells (AC) (Pellegatta *et al*, 2006).

Here, we used immunohistochemistry to assess GLAST expression in a large series of gliomas (n=144). Although in 28 out of 32 samples (88%) of low-grade gliomas (LGG, grade I and II, n=32) GLAST expression was negative (**Fig. 1A**, panel i) or only mildly positive (**Fig. 1A**, panel ii), we observed diffuse GLAST expression in 7 out of 9 anaplastic grade III astrocytomas (77%) (**Fig. 1A**, panel iii). In the GBM samples (n=103), however, there was diffuse positive staining (**Fig.1B**), with some specimens displaying intense GLAST expression (53 out of 103; 51%) and others showing lower levels of GLAST expression (50 out of 103; 49%). GLAST was mainly localized to the plasma membrane in the tumor cells.

**Figure 1.**
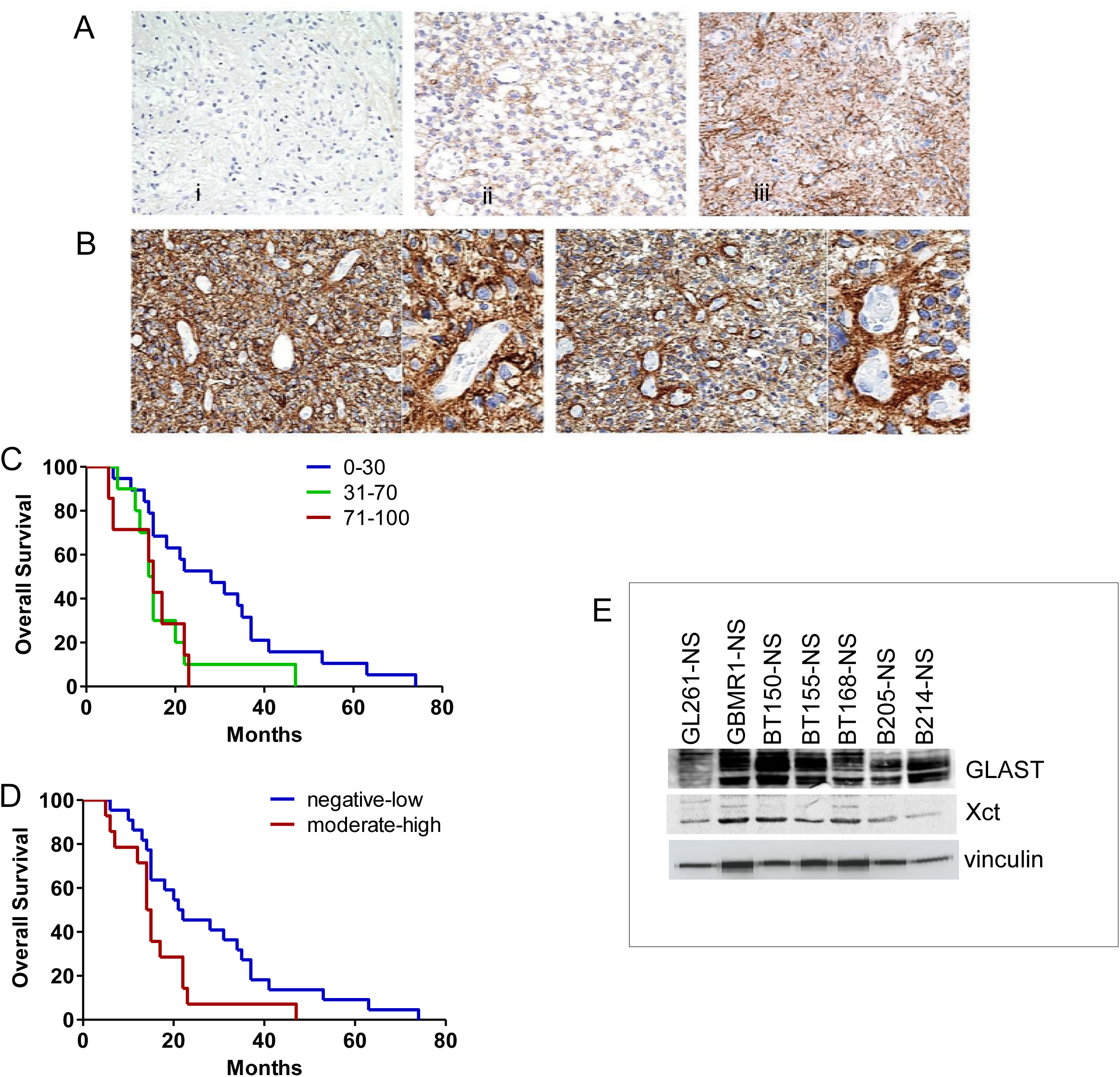
GLAST is expressed in glioblastoma specimens and correlates with patient survival. A, B. Representative examples of GLAST expression and immunoreactivity in i. grade I, ii. grade II (total n = 32), and iii. grade III glioma specimens (n = 9; 20X magnification) and glioblastomas (n = 103; 20X and 40X magnification). C, D. Kaplan-Meier survival curves showing a significant correlation between GLAST expression and poor prognosis (% GLAST^pos^ cells: *P* = 0.01 71-100 vs. 0-30; *P* = 0.04 31-70 vs 0-30; GLAST intensity *P* = 0.02 moderate-high vs negative-low by log-rank test; n=36 patients). E. Immunoblots showing GLAST and xCT protein expression in murine GL261 glioma and 6 human GSC lines, indicated as neurospheres (NS), normalized to vinculin. Densitometric quantification of GLAST and xCT expression in GL261-NS and representative GBM-NS.

After analyzing a group of patients (n=36, Table S1) who all received standard post-surgical concomitant radio-chemotherapy (Stupp *et al*, 2005, 2009), we found that the presence of a high percentage of GLAST-positive cells and strong or moderate reactivity significantly correlated with lower overall survival, supporting a potential role of GLAST as a prognostic marker for GBM (**Fig. 1, C and D**). We also investigated expression of the cystine/glutamate antiporter (xCT) in the GBM specimens. (**Fig. S1**). GLAST and xCT expression patterns were also evaluated by western blot analysis in GL261 cells as well as representative primary cell lines cultured in vitro as GBM neurospheres (GBM-NS). (**Fig. 1E**).

Taken together, these data indicate that GBM specimens and corresponding GSCs express GLAST at high levels and encourage further investigations into the functional consequences of GLAST expression.

### Impact of augmented and inhibited GLAST expression on tumor progression

To verify the involvement of GLAST in tumor progression, we isolated GLAST-expressing GL261 murine cells and human GSCs using magnetic cell sorting. The positive fraction showed a 10- to 14-fold enrichment in GLAST-expressing cells. Next, murine and human GLAST^high^ cells were injected intracranially, and they were significantly more aggressive than GLAST^low^ or unsorted cells, as defined by survival analysis (P < 0.001, **Fig. 2A, B**). Furthermore, magnetic resonance imaging (MRI) confirmed that GLAST^high^ and GLAST^low^ gliomas of murine (**Fig. 2C**) and human (**Fig. 2D**) origin showed different patterns of progression. Gliomas generated from GL261-GLAST^high^ cells infiltrated the contralateral hemisphere and disseminated in the third ventricle 21 days after intracranial injection (**Fig. 2C**, center). In contrast, gliomas generated from GLAST^low^ cells showed more demarcated borders 25 days after injection, indicating reduced invasive ability and less aggressiveness compared to gliomas from unsorted cells (**Fig. 2C**, right). Similar results were obtained when evaluating human GSCs. Twenty-seven days after implantation, gliomas were present in mice injected with GLAST^high^ human GSCs, whereas mice injected with GLAST^low^ GSCs had not developed tumors (**Fig. 2D**). On day 46 after injection, gliomas from GLAST^high^ GSCs showed much more rapid development than those from GLAST^low^ GSCs, where enhanced lesions were smaller and more circumscribed (**Fig. 2D**).

**Figure 2.**
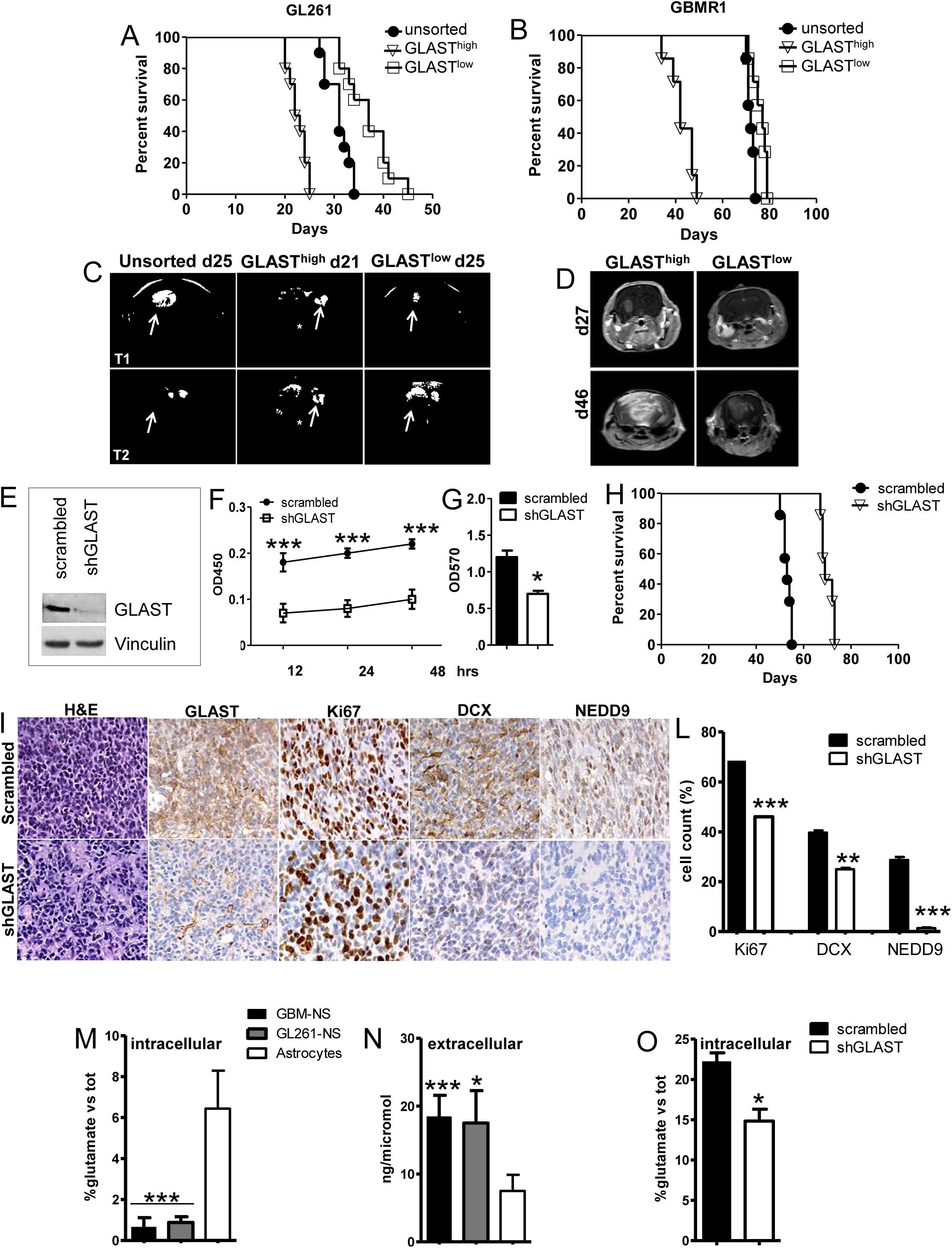
GLAST is involved in tumor invasion and glutamate release. A, B. Kaplan-Meier survival curves showing that murine and human GLAST^high^ cells are more aggressive than GLAST^low^ and unsorted cells (P<0.0001 by log-rank test; n=10 or 7 mice/group, respectively). GL261-GLAST^low^ glioma-bearing mice showed significantly shorter survival than those bearing gliomas from unsorted cells (P=0.0021 by log-rank test; n=10 mice/group, respectively). C, D. MRI performed on mice bearing murine and xenograft gliomas (n = 10/group, respectively). (C upper panels) T1-weighted images (T1-wi) with contrast medium injection. (C lower panels) T2-weighted images (T2-wi) showing extensive lesions infiltrating the contralateral hemisphere (arrow) and disseminating away from the tumor mass (asterisk) 21 days after GLAST^pos^ cell injection. Unsorted and GLAST^neg^ gliomas were less infiltrative on day 25 and preferentially developed in the homolateral hemisphere (arrow). D. T1-wi with contrast medium injection showing GLAST^high^ and GLAST^low^ xenograft models 27 days after tumor cell implantation (upper panels) as well as GLAST^pos^ and GLAST^neg^ xenograft models 46 days after tumor cell implantation (lower panels). E. GLAST-silencing efficacy in GCS was evaluated by western blotting. F. Cell proliferation was evaluated with a colorimetric method using WST reagent in both shGLAST and scrambled cells (Two-tailed Student’s t-test; ** *P*< 0.005; *** *P*< 0.0005; n=8 biologic replicates). G. Quantification of cell migration using Matrigel-coated transwell plates (Two-tailed Student’s t-test; *P*< 0.005 shGLAST vs. scrambled cells; n= 3). H. Kaplan-Meier survival curves showing that mice injected with shGLAST GSCs had shorter survival than those injected with scrambled GSCs (P<0.0001 by log-rank test; n=7 mice per group). I. Representative histological analysis of GLAST, Ki67, DCX and NEDD9 expression performed on serial sections of gliomas from mice injected with scrambled or shGLAST GSCs. L. Immunohistochemical quantification of Ki67-, DCX-, and NEDD9-positive cells in shGLAST vs. scrambled gliomas (Two-tailed Student’s t-test; ** P=0.001, *** *P* = 0.0001; n= 3). Data are presented as mean ± SD. M, N. Evaluation of glutamate uptake and release using a colorimetric assay performed on 15 GSC cell lines indicated as GBM-NS (uptake: 0.6±0.5% glutamate vs tot, release: 18.4.0±3.2 ng/microl) as well as on GL261-NS (uptake: 0.9±0.3% glutamate vs tot; release: 17.5±4.8 ng/microl) and normal astrocytes (uptake: 7.4±2.6% glutamate vs tot, release 7.5±2.4 ng/microl) (Two-tailed Student’s t-test; *** *P*< 0.0005; ** *P* < 0.01 vs astrocytes). Uptake and release assays were performed n = 3 times. Data are presented as mean ± SD. O. Detection of glutamate release from shGLAST vs scrambled GSCs (Two-tailed Student’s t-test; *P*< 0.01; n = 2 biological replicates). Data are presented as mean ± SD.

To evaluate the functional role of GLAST in human GSCs, we targeted GLAST expression using lentiviral particles expressing shRNA (shGLAST); as a negative control, we used lentiviral particles encoding a nonspecific shRNA (scrambled). Silencing efficiency was evaluated by immunoblotting (**Fig. 2E**). Using the shGLAST construct, the protein was silenced by nearly 90% compared with scrambled construct.

In vitro proliferation assays revealed significantly less proliferation of the shGLAST GSCs compared to the scrambled GSCs (**Fig. 2F**). The shGLAST cells also had significantly lower migration capacity (−2.0-fold vs. scrambled cells) (**Fig. 2G**). In vivo, down-regulation of GLAST inhibited tumor progression and improved the survival of nude mice injected intracranially with shGLAST GSCs (P<0.0001vs. scrambled GSCs) (**Fig. 2H**).

Histological analysis showed the almost complete absence of GLAST expression in the shGLAST glioma cells compared to the scrambled controls. These tumors also had a lower proliferation index based on the number of Ki67-positive cells. The cells also had reduced migration ability, which was assessed based on the identification of doublecortin (DCX)-positive cells (Daou *et al*, 2005), compared to those from the scrambled gliomas. Notably, the invasion marker NEDD-9 (Speranza *et al*, 2012) was totally absent from the shGLAST gliomas compared to the scrambled controls (**Fig. 2I, L**).

Because GLAST functions as a glutamate transporter, we next evaluated glutamate uptake in murine and human GSCs; murine astrocytes were used as a control. Glutamate uptake in human and murine neurospheres (GBM-NS and GL261-NS) was significantly lower than in normal astrocytes (**Fig. 2M**). In contrast, the extracellular level of glutamate in the medium of NS was higher compared to astrocytes (**Fig. 2N**). A significant reduction in extracellular glutamate was detected in vitro in shGLAST compared to scrambled GSCs (**Fig. 2O**), suggesting a direct relationship between GLAST expression and the passive movement out of the cells that we can consider as a glutamate release.

### GLAST-enriched gliomas show increased glutamate release

To examine whether GLAST expression increases glutamate release in vivo, we injected immunocompetent mice with GLAST^high^ (n=12) or GLAST^low^ GL261-NS cells (n=12) obtained by immunomagnetic sorting. Magnetic resonance spectroscopy (MRS) was used to quantify glutamate concentration (Doblas *et al*, 2012). The physiological concentration of glutamate measured in all mice before GL261-NS implantation was 7.2±1.3 mM±% Cramer-Rao bounds (CRB) error. Notably, for all examined GLAST^high^ gliomas, the glutamate concentration increased before tumor appearance at both the injection site (from 7.2±1.3 to 11.4±5.0 mM±%CRB) (**Fig. 3A**) and in the contralateral hemisphere (to 11.0±5.0 mM±%CRB) (**Fig. 3B**). The glutamate concentration was lower in GLAST^low^ gliomas, with the initial concentration being similar to physiological levels in both hemispheres (**Fig. 3C, D**). The increased levels of glutamate in the GLAST^high^ gliomas occurred prior to increase in tumor mass based on MRI evidence and were predictive of tumor cell infiltration. MRI performed at later time points confirmed that gliomas from GLAST^high^ cells grew in the two hemispheres as a consequence of early infiltration (**Fig. 2C, Fig. 3E**). In contrast, GLAST^low^ gliomas showed less infiltration and had more demarcated borders, indicating their reduced invasive ability (**Fig. 2C**).

**Figure 3.**
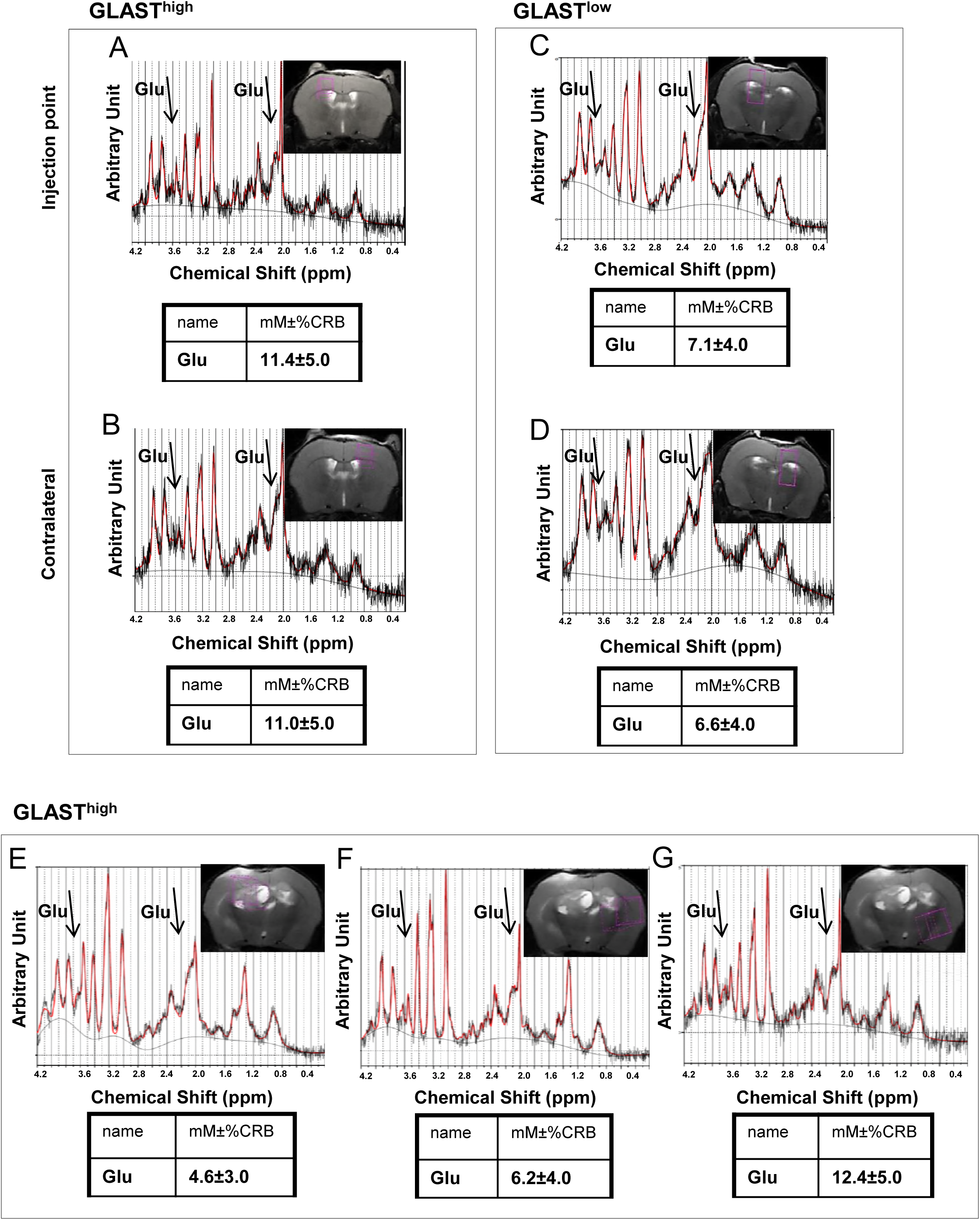
Glutamate concentration is higher in GLAST^high^ versus GLAST^low^ gliomas. A, B Variations in brain glutamate concentration measured at the injection site of GL261-NS cells (n = 12/group). C, D. Variations in brain glutamate concentration measured in the contralateral hemisphere in GLAST^low^ gliomas (n = 12/group). 1H-MRS spectra were obtained from single voxels (∼10 µL) on T2-weighted images located inside a tumor, at the injection site, one week after tumor implantation, and symmetrically in the contralateral hemisphere (purple rectangle) as well as in brain regions not yet exhibiting detectable solid tumor. The absolute concentrations of Glutamate metabolite (arrows highlites Glu MRS peaks) computed by LcModel software (red line) are listed under the corresponding spectra. E-G. MRS was performed three weeks after tumor implantation on lesions from GLAST^high^ GL261 cells infiltrating the contralateral hemisphere and revealed reduced glutamate levels in the tumor mass at the injection point (**E**) as well as in the contralateral hemisphere (**F**) but increased glutamate levels in the peritumoral region (**G**). The metabolite concentrations (mM±%CRB) are representative of n = 12 mice/group.

We also evaluated glutamate concentrations in mice with GLAST^high^ tumors that had invaded the contralateral hemisphere. Decreased glutamate concentration was found within the tumor mass (**Fig. 3E**), with a slight increase in the contralateral hemisphere (**Fig. 3F**). The highest levels of glutamate were measured in peritumoral regions contralaterally (**Fig. 3G**). Overall, the above results support a key role of glutamate in tumor cell infiltration into the opposite hemisphere.

### GSCs show reduced glutamate uptake due to their lack of Na/K-ATPase expression

As discussed above, GLAST is responsible for glutamate uptake, and its activity depends on the Na+ electrochemical gradient generated by Na^+^/K^+^-ATPase (Attwell *et al*, 1993; Rose *et al*, 2009; Szatkowski *et al*, 1990). Therefore, we next evaluated Na^+^/K^+^-ATPase expression in GBM specimens and GSCs. Real-time PCR was performed on 29 GBM specimens and showed that expression of the ATP1A1, ATP1A2 and ATP1B2 genes, significantly decreased compared to that measured in normal brain (NB, *P*<0.005) Specifically, we analyzed the α2 isoform, which is primarily expressed in adult brain astrocytes (McGrail *et al*, 1991), and α1, to exclude any mechanism of compensation Alpha2 is typically combined with the β2 subunit (Cameron *et al*, 1994), which has been implicated in regulating both the activity and conformational stability of the α subunit (Chow & Forte, 1995). (**Fig. 4A**). Immunoblotting confirmed that α2, β2 and α1 were either absent or barely expressed in all GSCs analyzed (**Fig. 4B and Fig. S1**). We further evaluated 12 GBM specimens that strongly expressed GLAST and found that few α1 and α2-positive cells were located near blood vessels and that these cells were virtually absent from tumor masses that exhibited very high levels of GLAST expression (Fig S1 and **Fig. 4C**). Interestingly, the α2 subunit co-localized with the endothelial marker CD31 (**Fig. 4D**). The down-regulation of α1 and α2 subunit associated with high expression of GLAST was confirmed by western blot performed on three different GBM representative for GLAST-high, intermediate and low expression (Fig S1D). To demonstrate that absence of the ATPase was responsible for the reduced glutamate uptake in GBM cells, we over-expressed the α2 and β2 subunits in a representative human GBM-NS (BT168-NS). We found that ATP1A2 and ATP1B2 were overexpressed compared to the control (MOCK) based on results from both Real Time (RT)-PCR and western blotting (**Fig. 4E** and **4F, respectively**). Furthermore, GBM-NS that expressed Na+/K+-ATPase (ATP1α2β2-NS) showed a restored ability to uptake glutamate compared to the control. However, treatment of ATP1α2β2-NS with UCPH-101, a selective GLAST inhibitor, significantly reduced glutamate uptake back to baseline (**Fig. 4G**). The increase in glutamate uptake induced apoptosis in the ATP1α2β2-NS based on flow cytometry results (**Fig. 4H, I**). Apoptotic cells were identified based on annexin V/propidium (LA=apoptosis) and caspase 3-7/SYTOX double-positive staining, and the proportion significantly increased in the ATP1α2β2-NS compared to the control. Furthermore, treatment of ATP1α2β2-NS with UCPH-101 reduced the percentage of apoptotic cells compared with no treatment.

**Figure 4.**
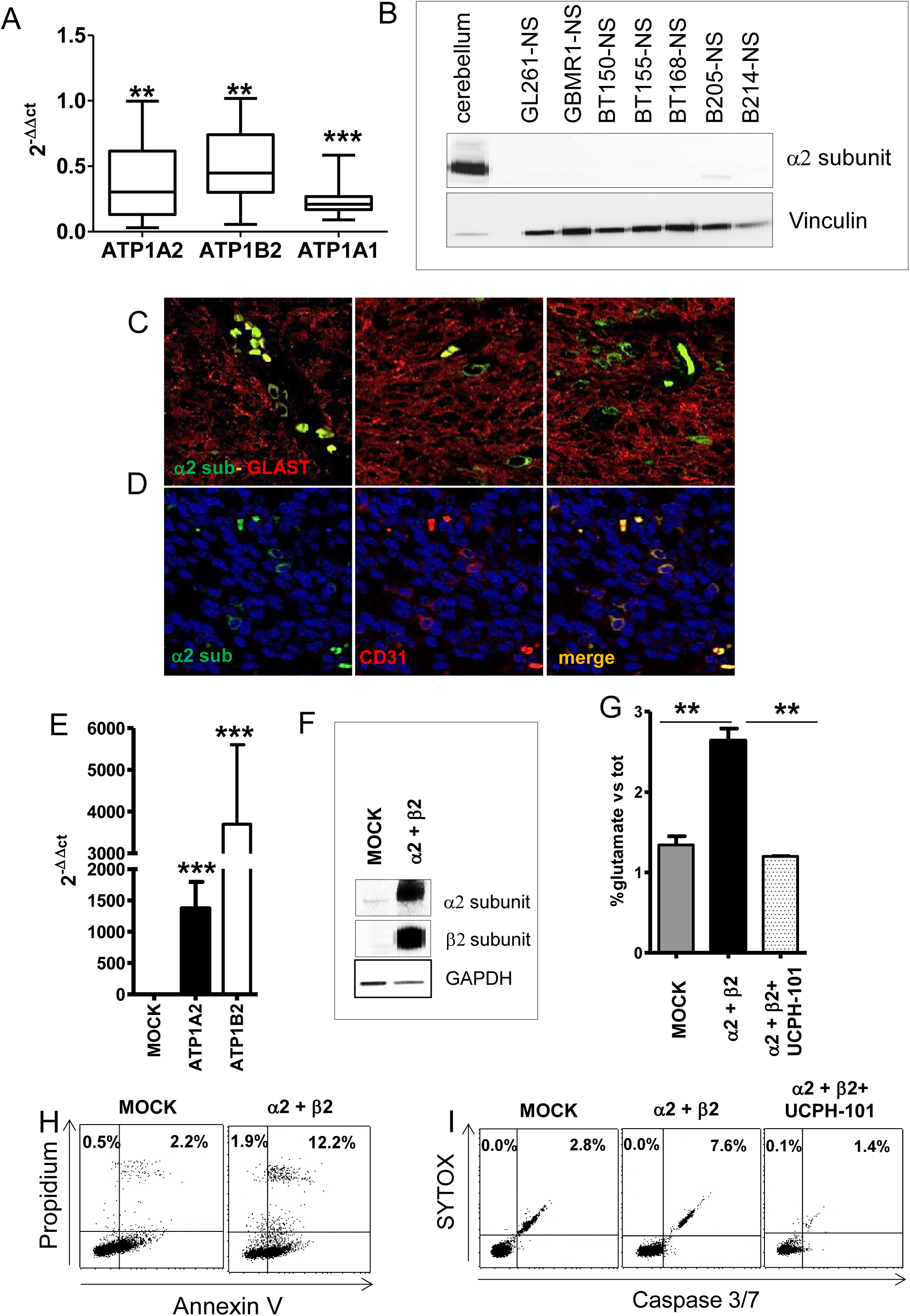
GSCs and glioblastoma specimens do not express Na^+^/K^+^-ATPase. A. Expression of ATP1A2, ATP1B2, ATP1A1 subunits evaluated by real-time PCR (Two-tailed Student’s t-test; *P*< 0.005 vs normal brain (2^-ΔΔCt^ =1); n= 29 GBM specimens. All data are presented as mean ± SD. B. Western blotting to detect α2 expression in GL261-GSCs and human GSC lines. C, D. Representative confocal analysis of immunofluorescence staining to detect expression of the α2 subunit and GLAST and the α2 subunit and CD31 in 12 human GBM specimens. E. Relative expression of the ATP1A2 and ATP1B2 subunits in transfected GSCs detected by real-time PCR. ATP1A2 and ATP1B2 expression was significantly higher in transfected cells than control cells (MOCK) (Two-tailed Student’s t-test; ***P<0.0001; n=3 biologic replicates). All data are presented as mean ± SD. F. Western blot showing overexpression of the α2 and β2 subunits in a transfected primary cell line. Both subunits showed significantly increased expression compared with the MOCK control. G. Intracellular glutamate levels in a primary cell line expressing the α2 and β2 subunits were evaluated using a colorimetric assay. The results showed significantly higher intracellular glutamate levels relative to the MOCK control, and this trend was abolished after UCPH-101 treatment (Two-tailed Student’s t-test; **P<0.001). H, I. Flow cytometry was used to evaluate apoptosis based on annexin V/propidium (H) and caspase 3-7/SYTOX staining (I). Cells expressing the α2 and β2 subunits showed a higher percentage of apoptosis based on annexin V/propidium double-positive staining (12.2%) as well as increased necrosis based on caspase3-7/SYTOX double-positive staining (7.6%) compared to the MOCK control. UCPH-101 treatment did not affect the viability of the transfected cells relative to the control cells.

Overall these data indicate that the absence of Na+/K+-ATPase on glioma membranes is responsible for the reversal in GLAST function.

### The GLAST inhibitor UCPH-101 causes apoptosis of GSCs and increases survival in vivo

To further assess the effects caused by specific inhibition of GLAST using UCPH-101, we treated GL261-NS, GBM-NS and astrocytes with 10 μM UCPH-101 or DMSO as a control. The GL261-NS and GBM-NS showed a significant increase in apoptosis based on annexin V/propidium- and caspase 3/7/SYTOX-positive staining after treatment with UCPH-101 (**Fig. 5 A, B, D**), whereas astrocytes were not affected by the treatment (**Fig. 5 C, D**). The percentages of both apoptotic cells (annexin V/propidium double-positive staining) and necrotic cells (caspase 3-7/SYTOX double-positive staining) significantly increased during the treatment.

**Figure 5.**
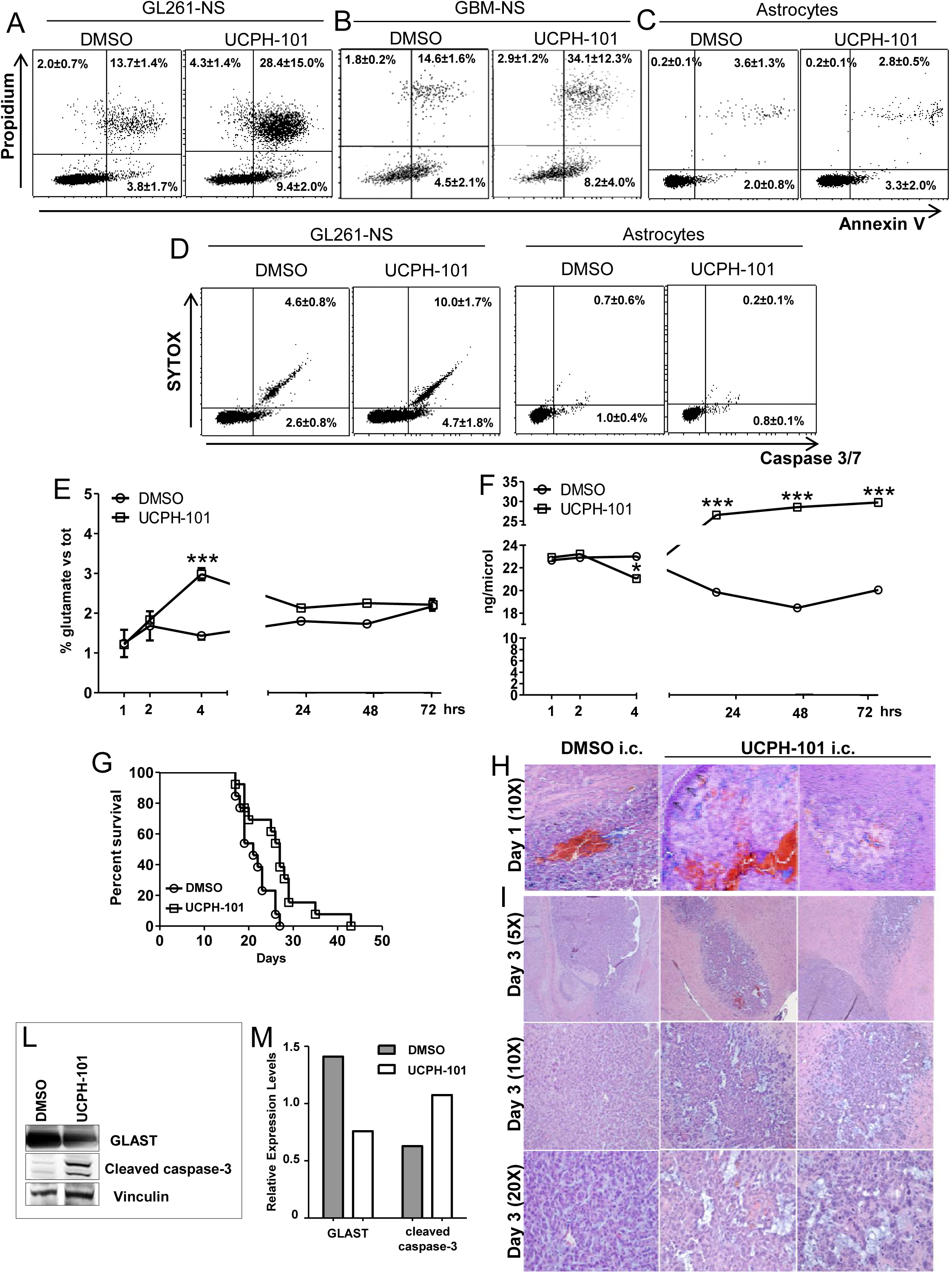
The GLAST inhibitor UCPH-101 caused GL261 cell apoptosis both in vitro and in vivo. A-D. Representative dot plots showing apoptosis evaluation by flow cytometry using annexin V/propidium staining in GL261-NS, GBM-NS and astrocytes (**A-C**) and caspase 3-7/SYTOX staining in GL261-NS and astrocytes (**D**). A. GL261-NS treated with UCPH-101 (10 μM) showed a higher percentage of late apoptotic cells based on annexin V/propidium double-positive staining (28.4±15% vs 13.7±1.4%, P=0.02) and of early apoptotic cells based on annexin V positive/propidium negative staining (9.4±2% vs 3.8±1.7%, P=0.01) compared to DMSO-treated cells. Data are presented as mean ± SD B. GBM-NS treated with UCPH-101 (10 μM) showed a higher percentage of late apoptotic cells based on annexin V/propidium double-positive staining (34.1±12.3% vs 14.6±1.6%, P<0.005) and of early apoptotic cells based on annexin V positive/propidium negative staining (8.2±4.0% vs 4.5±2.1%, P=0.04) compared to DMSO-treated cells. C. Astrocytes treated with UCPH-101 (10 μM) showed basal levels of apoptosis. D. GL261-NS treated with UCPH-101 (10 μM) showed a significant increase in necrotic cells based on caspase 3-7/SYTOX double-positive staining (10.0±1.7% vs 4.6±0.8%) and of apoptotic cells based on caspase3-7 positive/SYTOX negative staining (2.6±0.8% vs 4.7±1.8%) relative to DMSO-treated cells. Astrocytes treated with UCPH-101 (10 μM) did not show necrosis (Two-tailed Student’s t-test, p < 0.01). Data are presented as mean ± SD of three independent experiments at the three different time points. E. Intracellular glutamate levels were evaluated in GL261-NS treated with UCPH-101 (10 μM) using a colorimetric assay. Four hours after UCPH-101 treatment, the GL261-NS showed significantly higher intracellular glutamate levels relative to DMSO-treated cells (Two-tailed Student’s t-test; ***P<0.0001; the uptake assay was repeated n= 3 time. Data are presented as mean ± SD). There was no significant difference in glutamate intracellular levels at later time points (24, 48 and 72 hrs). F. Extracellular glutamate levels were evaluated in GL261-NS treated with UCPH-101 (10 μM) using a colorimetric assay. Extracellular glutamate levels significantly increased in the GL261-NS starting after 4 hrs of UCPH-101 treatment and this effect persisted at later time points (24, 48 and 72 hrs) relative to DMSO-treated cells (Two-tailed Student’s t-test; *P<0.01, ***P<0.0001; the release assay was repeated n = 3 time. Data are presented as mean ± SD). G. Kaplan-Meier survival curves showing that a single intratumoral delivery of UCPH-101 (10 nmol/μl) 9 days after GL261-NS implantation markedly improved survival (P=0.0075 by log-rank test; n= 7mice/group). H. H&E of GL261-gliomas one day after DMSO or UCPH-101 administration (10 nmol/μl i.c.). Representative gliomas treated with DMSO showed high cellularity and some damaged areas due to vehicle administration, and a ring of tumor cells appeared to migrate to the borders of the damaged areas (black arrows). I. H&E of GL261-gliomas three days after DMSO or UCPH-101 administration (10 nmol/μl i.c.). The control glioma showed a larger mass than the UCPH-101-treated glioma (upper panel magnification, 5X). The center and lower panels show 10X and 20X magnification, respectively. A representative control glioma showed higher cellularity compared with the treated gliomas, which are characterized by several necrotic areas. Pictures are representative of n = 4 per group per time points. L. Western blot showing GLAST and cleaved caspase-3 expression in GL261-gliomas treated with UCPH-101 (10 nmol/μl). The treated GL261-gliomas showed decreased GLAST expression and increased cleaved caspase-3 expression relative to DMSO-treated gliomas. Vinculin was used as a loading control. The immunoblot is representative of n = 2 explanted glioma/group. M. Densitometric quantification of GLAST and cleaved caspase-3 expression in GL261-tumors treated with UCPH-101.

The effects of UCPH-101 on glutamate uptake and release showed a time-dependent behavior (**Fig. 5E, F**). GL261-NS showed increased intracellular levels of glutamate at early time points (1–4 hours) after treatment with UCPH-101, but this difference was lost at later time points, and extracellular glutamate significantly decreased 4 hours after the treatment (**Fig. 5E**). Consistently, extracellular levels of glutamate dropped four hours after addition of the GLAST inhibitor UCPH-101. At later time points, there was a dramatic reversal of this trend, and a significant increase of extracellular glutamate accumulating in the cell culture medium (**Fig. 5F**) in parallel to cell death and the subsequent passive release of glutamate into the medium.

We also examined the effects of UCPH-101 treatment in vivo by administering a single dose of the inhibitor into GL261 glioma-bearing mice via intra-tumoral injection 9 days after tumor implantation. This single injection was sufficient to significantly increase animal survival (p=0.0075, **Fig. 5G**) without causing toxicity. In mice sacrificed one day after treatment, gliomas from vehicle-treated controls showed either slight or no effects (**Fig. 5H** left), whereas UCPH-101-treated mice showed substantial disruptions in glioma architecture highlighted by large necrotic areas (**Fig. 5H**, right)) surrounded by residual tumor cells (**Fig. 5H**, arrows). Three days after the administration, control gliomas were larger than UCPH-101-treated gliomas and characterized by high cellularity (**Fig. 5I**). Necrotic areas were still evident in the UCPH-treated gliomas, although other areas showed some recovery of tumor growth. We next investigated the effects of the treatment on GLAST expression in explanted tumors. UCPH-101 caused an early decrease in GLAST expression based on the results of western blot analysis (**Fig. 5L, M**). Furthermore, gliomas explanted from the UCPH-101-treated mice showed a marked increase in cleaved caspase-3 expression.

### GLAST expression is up-regulated in response to STAT3 activation by glutamate

To investigate the signaling mechanism connecting glutamate and GLAST up-regulation, we assessed the relationship between STAT3 and GLAST expression, as the STAT pathway has a role in GLAST expression in astrocytes (Wang *et al*, 2016). Glutamate concentrations were measured by MRS in two different regions of GL261-GLAST^high^ gliomas.

Higher levels of glutamate were found in the peritumoral region rather than within the tumor mass (**Fig. 6A**). We then separately examined the tumor mass and peritumoral region in explanted tumor and found that both GLAST expression and STAT3 activation, evaluated based on the presence of STAT3 phosphorylated at Tyr-705, were lost in the tumor mass, where the glutamate concentration was lower than physiological levels. In contrast, glutamate levels were higher than physiological levels in the peritumoral areas and, accordingly, there were high levels of GLAST expression and STAT3 activation (**Fig. 6B**).

**Figure 6.**
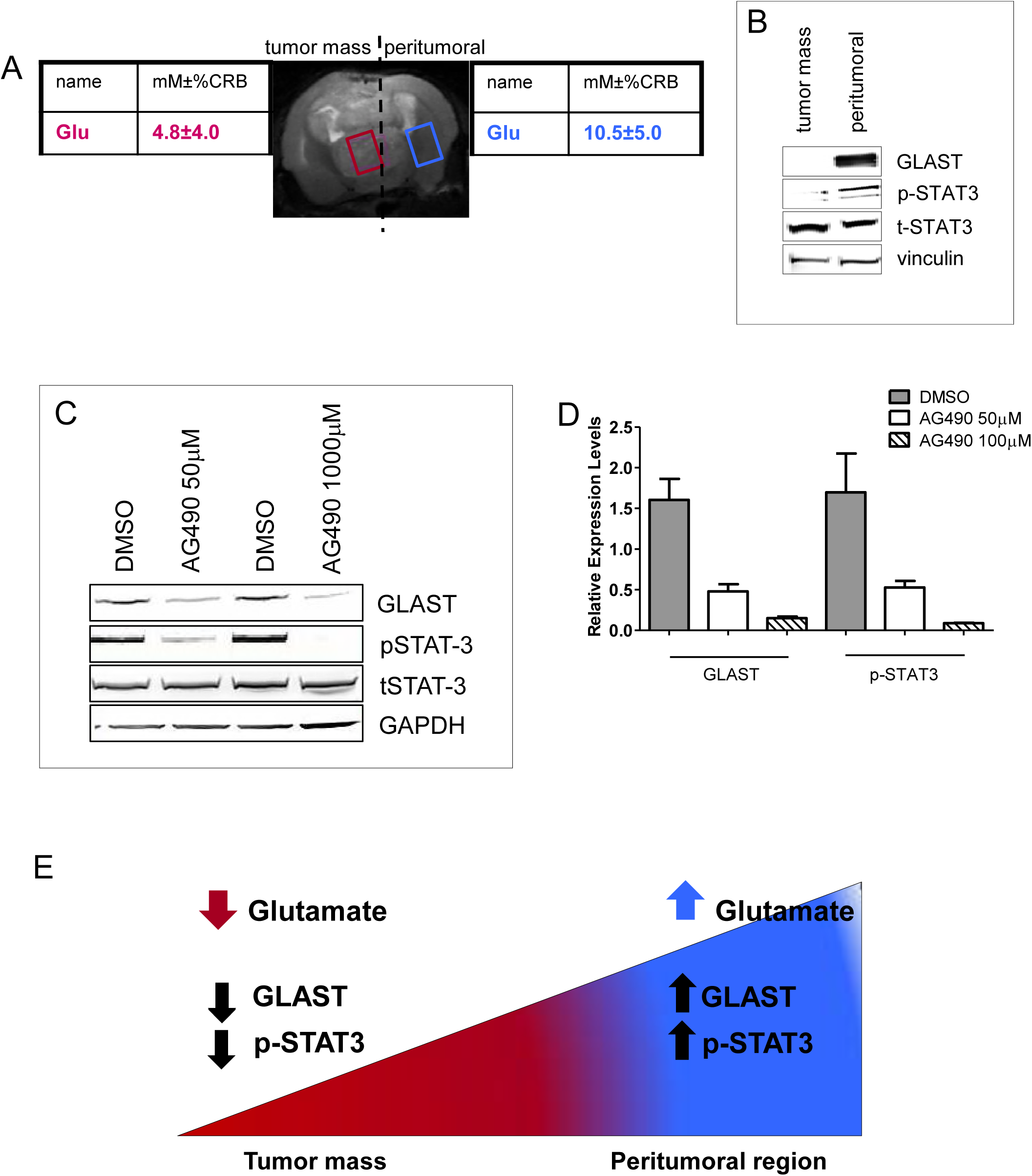
STAT3 activation regulates GLAST expression. A. MRS performed on GL261 GLAST+ gliomas 25 days after tumor implantation showed that glutamate levels decreased in the tumor center (4.8±4.0mM±%Cramer-Rao bounds CRB error, purple rectangle) with no evidence of necrosis. Glutamate levels were higher than the physiological concentration at the borders of the tumor in the contralateral hemisphere (peritumoral area, 10.5±5.0 mM±%CRB, blue rectangle). The metabolite concentrations (mM±%CRB) are representative of n = 6 mice/group. B. Separate immunoblotting of explanted gliomas at the tumor center and peritumoral area showed a loss of GLAST expression in the tumor center, where the glutamate concentration was very low and STAT3 was inactivated. The opposite trend was observed at the tumor periphery, where increased glutamate levels corresponded to increased GLAST expression and STAT3 activation (i.e., phosphorylation of Tyr705). C. Western blot performed on GL261-NS showing that AG490, a specific STAT3 inhibitor, decreases GLAST expression in a dose-dependent manner. GAPDH was used as a loading control. The experiment was performed twice. D. Densitometric quantification of GLAST and pSTAT-3 expression in GL261 tumors treated with DMSO or AG490 at two different doses. E. Schematic representing glutamate levels in the tumor periphery compared to the tumor center and the resultant influence on GLAST expression. Glutamate can up-regulate GLAST expression via STAT3 activation (pSTAT3).

To further confirm the involvement of STAT3 activation in regulating GLAST expression, we inhibited the JAK/STAT3 pathway in GL261-NS cells using AG490, a tyrphostin tyrosine kinase inhibitor. To accomplish this, GL261-NS cells were treated with 50 μM or 100 μM AG490 or DMSO for 24 hours. Using immunoblotting, we observed significant reductions in GLAST and pSTAT3 Tyr 705 expression in NS treated with AG490 compared with DMSO (**Fig. 6C, D**). Our results suggest that glutamate may play a role in triggering up-regulation of GLAST expression through STAT3 activation (pSTAT3) and that glutamate levels in peritumoral regions may influence the expression of GLAST as exemplified in **Fig. 6E**.

## Discussion

Here, we demonstrated that GLAST, an astrocyte transporter responsible for glutamate uptake and a radial glia marker (Hartfuss *et al*, 2001; Merkle *et al*, 2004), is expressed on GBM cell surfaces and helps these cells migrate towards and invade normal surrounding tissue. In a previous study, we also showed that GLAST is highly expressed in GL261 GSCs (Pellegatta *et al*, 2006) and more recently, we showed that injection of GLAST peptides into mice promotes specific anti-tumor responses towards murine gliomas (Cantini *et al*, 2012).

Glutamate metabolism and trafficking have been shown to be profoundly altered in numerous cell and animal models of gliomas as well as in human glioma patients (Lyons *et al*, 2007; Takano *et al*, 2001; Ye & Sontheimer, 1999b). The cystine-glutamate exchanger xCT that is instrumental in GBM release of excitotoxic concentrations of glutamate (Takeuchi *et al*, 2013) is expressed in 50% of GBMs and has been related to the outcome of patients with GBM (Robert *et al*, 2015). Here, we observed that GLAST is highly expressed on the plasma membrane in GBM cells. Interestingly, GBMs with no or low expression of GLAST showed xCT localization to the cell surface. GSCs grown in culture demonstrated impaired glutamate uptake and a predisposition to glutamate release. Accordingly, inhibition of GLAST expression reduced glutamate release in vitro and limited tumor invasion and progression in GBM xenografts. It has previously been shown that peritumoral glutamate can contribute to brain edema (Savaskan *et al*, 2008), which amplifies the tumor mass effect and is often responsible for symptoms related to gliomas, including seizures (Buckingham & Robel, 2013; Buckingham *et al*, 2011; Yuen *et al*, 2012). Notably, MRS revealed higher concentrations of glutamate in mice injected with GLAST-enriched cells before the appearance of tumor mass, with a robust increase in glutamate in peritumoral areas where GLAST was expressed.

Increased GLAST expression was associated with weak/absent expression of Na+/K+-ATPase, which maintains the electrochemical gradient necessary for glutamate uptake. The collapse of the Na+ gradient under pathological conditions, including ischemia, has been shown to be responsible for uptake reversal and consequent release of glutamate (Szatkowski *et al*, 1990). Glutamate release by astrocytes is rapid and reversible and occurs through a complex mechanism that involves local energy failure, inhibition of glycolysis, and depolarization of the cell membrane (Hertz, 1977; Kimelberg *et al*, 1990; Nicholls & Attwell, 1990; Ogata *et al*, 1995).

Overall, we found that GBM cells express GLAST but not Na+/K+-ATPase. Previous reports have indicated that down-regulation or aberrant expression of Na+/K+-ATPase subunits has a role in the development and progression of various cancer types (Mijatovic *et al*, 2007). The β2-subunit, in particular, is reported to be completely absent from glioma cells (Senner *et al*, 2003) or down-regulated in GBMs in association with increased invasion (Senner *et al*, 2003; Sun *et al*, 2013). The α1 was found expressed in GBM and in U373-MG established cells, in contrast with our observation (Lefranc *et al*, 2008). The α2 subunit has been reported to be strongly co-expressed with GLAST in cerebellar Bergmann glia, in astrocytes and in the cerebral cortex (Rose *et al*, 2009; Illarionova *et al*, 2014). Here, we showed that inhibition of GLAST expression reduced glutamate release in vitro and limited tumor invasion and progression in xenograft models. Therefore, glutamate is crucial in promoting GBM infiltration and long-distance diffusion into normal brain tissue.

Our data unveil a novel relationship between glutamate release and glioma progression. The inverse correlation that exists between GLAST expression and patient survival suggests that this mechanism is clinically relevant. Thus, therapeutic targeting of GLAST by immunotherapy, which we previously reported (Cantini *et al*, 2012), or through the use of GLAST inhibitors may help control invasion and edema during GBM evolution.

We also evaluated the use of the GLAST inhibitor UCPH-101 in gliomas for the first time (Abrahamsen *et al*, 2013). Intriguingly, UCPH-101 induced cell death in GBM-GSCs, but not in astrocytes. Based on these results, it appears that glutamate is released by GBM-GSCs through GLAST. We speculate that inhibition of this release increases intracellular glutamate levels in GBM-GSCs, but not in astrocytes, and that these levels eventually become toxic and induce apoptosis.

Previous data have shown that UCPH-101 has a limited ability to penetrate into the brain (Erichsen *et al*, 2010). Even though the BBB has been shown to be disrupted in the presence of GL261 gliomas (Leten *et al*, 2014), here we treated glioma-bearing mice via intratumoral administration. We found that a single injection of UCPH-101 was sufficient to prolong the survival of glioma-bearing mice and induce apoptosis of tumor cells in vitro. These results indicate the relevance of perturbed glutamate trafficking in glioma evolution. Additionally, we found that the STAT3 signaling pathway is triggered by glutamate and responsible for GLAST up-regulation. STAT3 expression is found in normal brain tissue, specifically in neurons, glial cells and Bergmann glia (Aguilera & Ortega, 1999; Park *et al*, 2012; Planas *et al*, 1997). Activation of STAT3 has been studied in detail and has an important role in the signaling cascade activated by glutamate (Aguilera & Ortega, 1999). In particular, it has been shown to protect against glutamate-induced neuronal death and excitotoxicity (Park *et al*, 2012). Astrocytes under chronic hypoxia showed decreased GLAST expression and function following STAT3 inactivation (i.e., after levels of pSTAT3-Tyr705 decreased). Conversely, up-regulation of GLAST expression has been demonstrated as an important indirect anti-apoptotic mechanism in the adult CNS (Koeberle & Bähr, 2008). Both astrocytes and GBM cells utilize STAT3 signaling to prevent cell death. Based on our data showing that GLAST expression decreased in glioma-bearing mice treated with UCPH-101, we hypothesize that the high levels of glutamate found in the tumor periphery induce GLAST up-regulation through STAT3 activation, and vice versa. Overall, our findings combined with the selective effect of UCPH-101 on glioma cells (reducing toxicity to normal brain) support that clinical testing of GLAST inhibitors should be performed as a potential treatment option for glioma.

## Material and Methods

### Tumor specimens and cell culture

Glioblastomas (GBM) and low grade gliomas (LGG) studied for GLAST quantification and survival analysis were obtained from Unit of Pathology, University of Brescia. All patients provided their informed consent. Primary cell lines were derived from GBM specimens obtained from the Department of Neurosurgery at the Istituto Neurologico Carlo Besta, according to a protocol approved by the institutional Ethical Committee. After GBM processing cell suspensions were cultured as neurospheres and defined GBM stem-like cells (GSCs) as previously described (Finocchiaro & Pellegatta, 2016), and cultured in standard medium containing B27 supplement and the mitogenic factors epidermal growth factor (EGF) and fibroblast growth factor b (bFGF). GL261 were grown as neurospheres (GL261-NS) using the same medium. Frozen specimens and GSCs are stored in the the Besta Brain Tumor Biobank (BBTB) established with the collaboration of a GLP cryo-management service provides by SOL (SOL Group Spa, Italy). All of the GSCs are tested for mycoplasma contamination by Promokine PCR Test Kit (PK-CA91).

### Magnetic sorting

The isolation of murine and human GSCs was performed using the anti-GLAST (ACSA-1) Microbead kit (Miltenyi Biotec#130-095-826) according to the manufacturer’s instruction. Briefly, dissociated neurospheres were labelled with anti-GLAST-Biotin and were subsequently incubated with Anti-Biotin MicroBeads. The purity of the enriched cells was verified using anti-GLAST antibody-PE (Miltenyi Biotec #130-095-82, 1:10) by MACSQuant analyzer (Miltenyi Biotec). Dead cells were excluded from the analysis based on their scatter and DAPI signals.

### Silencing of GLAST expression

GSCs were transduced with lentiviral particles (MISSION shRNA Lentiviral Vectors, Sigma Aldrich) containing GLAST specific shRNA sequences (shGLAST) according to the manufacturer’s recommendations. Four different GLAST-specific sequences were screened, and the most efficient sequence was chosen (TRCN0000043195). As a negative control, we used shRNA Lentiviral Particles encoding non-specific shRNA (scrambled cells). Four days after infection, cells were selected for puromycin resistance (1.5 μg/ml) for at least one week.

### In vivo experiments

Five weeks-old female C57BL/6N (n=10/group for survival studies, figure 2a) and CD1 nude (nude CD1 (HO): CD1-Foxn1nu) mice (n=7/group for survival studies, figure 2b) were intracranially injected with 1 × 10^5^GL261-NS or GSCs respectively, in 2microL PBS1X. The coordinates, with respect to the bregma, were 0.7 mm post, 3 mm left lateral, 3.5 mm deep, and within the nucleus caudatum. Five weeks-old female CD1 nude mice (n=7/group for survival, figure 2h; n=3/group for histological studies; figure 2i) were intracranially injected with 1 × 10^5^ scrambled or shGLAST GSCs. Five weeks-old female C57BL/6N mice (n=7/group for survival analysis, figure 5g; n=4 for histological studies; figure 5h-5i) were injected with 1 × 10^5^ GL261-NS and treated with 10nmol/μl of UCPH-101 (Tocris Bioscience) or DMSO as negative control, nine days after tumor implantation. For survival studies mice were monitored every day and when suffering euthanized; for histological studies mice brains were collected and fixed in 4% paraformaldehyde. We placed no more than 5 mice per cage and we used a 12 light/12 dark cycle.

All mice were purchased from Charles River Laboratories (Calco, Italy) and all the experiments were performed in accordance with the directives of Istituto Neurologico C. Besta and Italian Ministry of Health. Animal experiments were performed in accordance to the Italian Principle of Laboratory Animal Care (D. Lgs. 26/2014) and European Communities Council Directives (86/609/EEC and 2010/63/UE).

### Na+/K+-ATPase overexpression in primary cell line

Primary cell line was seeded in 6-well plates at 2×10^6 cells/well 24hrs prior to the transfection. Cells were transfected with 5μg of ATP1B2-DNA vector (OriGene), ATP1A2-DNA vector (OriGene) or both using Lipofectamine 2000 Transfection Reagent (Life Technologies), according to manufacturer’s instruction. Twenty-four hours after transfection, cells were split and grown in fresh medium containing 0.5mg/ml of G418 (Lonza) as selective agent.

### Glutamate detection, proliferation, migration and apoptosis assays

Murine and human GSCs cells were cultured in DMEM + Ham’s F-12 without L-glutamine (PAA Laboratories-GE Healthcare). Glutamate quantification was performed using a colorimetric assay (Biovision) according to the manufacturer’s instructions. Release and uptake of glutamate were evaluated by testing the cell supernatants and cells homogenized in assay buffer, respectively. Normalization was obtained using the microBCA kit (Thermo Scientific). The proliferation kinetics were measured by plating 8000 cells/ cm^2^ in 25cm^2^-flasks. Cells were counted using the trypan blue exclusion test, which was performed every 3 days. Cells were seeded at 2000 cells/well, and cell proliferation was measured at different time points after plating using the cell proliferation reagent WST-1 (Roche Applied Science). Eight replicates per point were completed. The absorbance was determined at 450 nm. Migration was assayed *in vitro* using the Transwell-96 system (Becton Dickinson) as recommended by the manufacturer. After 24 and 48 h, the migrating cells were stained with crystal violet solubilised in 10% acetic acid, and the absorbance was determined at 570 nm. GSCs were cultured in DMEM + Ham’s F12 without L-glutamine (PAA Laboratories-GE Healthcare) and murine astrocytes were used as control. Cells were seeded at 2×10^5^ cells/well and treated with 10 μM of UCPH-101 or vehicle (DMSO). Cells were collected at different time point and apoptosis was evaluated using the Annexin V-FITC kit (Miltenyi Biotec) and the CellEvent™ Caspase-3/7 Green Flow Cytometry Assay Kit (Life Technologies) according to the manufacturer’s instructions.

### Immunoblot analysis and antibodies

The extraction of protein from cell cultures or freshly explanted tumors was made in lysis buffer (PBS containing 0.5% sodium deoxycholate, 1% NP40, 0.1% SDS, 1 mM sodium orthovanadate, 1 mM PMSF, 2mM NaF, 1 mg/ml of leupeptin and Aprotinin), protein concentrations were assessed using MicroBCA protein assay kit (ThermoFisher Scientific, Inc.), the samples were resolved on a 10% SDS-PAGE gel, and electroblotted onto nitrocellulose membranes. Non-specific protein binding-sites on membranes were blocked by incubation for 3 hours in 5% BSA in TBS-T before incubating with primary antibodies, at recommended dilutions, for 2–16 hours at 4°C. The membranes were incubated with the following primary antibodies diluted in blocking solution: anti-GLAST (1:200, Santa Cruz Biotechnology #sc-7757), anti-Na+/K+-ATPase α1, α2 (1:1000, Millipore #05-369 and #07-674) and β2 (1:1000, Thermo Fisher Scientific, Inc. #PA5-26279), anti-Vinculin (1:5000, Santa Cruz #sc-25336) and anti-GAPDH (1:3000, Sigma Aldrich #G8795), anti-Cleaved Caspase 3 (Asp175) (1:1000, Cell Signaling Technology #9661), anti-STAT3 and anti-pSTAT3 (Tyr-705) (1:1000, Cell Signaling Technology #9132 and #9131). The primary antibody interaction was followed by incubation with a secondary HRP-conjugated antibody diluted in blocking solution, anti-rabbit, anti-mouse or anti-goat (1:3000, Bethyl Laboratories, Inc.#A120-101P, #A90-117P and #A50-100P). Immunoreactive species were detected by chemiluminescence reaction using the ECL (enhanced chemiluminescence) Plus kit (Amersham, GE Healthcare). All antibody validation information is available in the product’s manual.

### RNA extraction and Real-Time PCR

Total RNA was extracted using Trizol reagent (Life Technologies) from human snap frozen tissues and GSCs. Total RNA was reverse-transcribed using a High Capacity cDNA Synthesis KIT (Applied Biosystems-Life Technologies). The expression of ATP1A1, ATP1A2 and ATP1B2 subunits were detected by SybrGreen chemistry performed on ViiA7 Real Time PCR system (Life Technologies), and normalized relative to beta-2 microglobulin. The RNA from non-transfected cells (MOCK) was used as the calibrator for the calculation of fold expression levels with the ΔΔCt method.

Oligo Sequences:

ATP1A1 Fw: ACGGCTTCCTCCCAATTCAC, Rv: CTGCTCATAGGTCCACTGCT; ATP1A2 Fw: TGCATTGAGCTCTCCTGTGG, Rv: CGTGGATAGACAGCTGGTACTT; ATP1B2 Fw: TTCCTCACCGCCATGTTCAC, Rv: GAATCATCAAGCCCGGTGTG; Beta-2-microglobulin Fw:GTGCTCGCGCTACTCTCTCT, Rv:CCCAGACACATAGCAATTCAG

### Immunohistochemical analyses and immunofluorescence

For IHC analyses human GBM and xenograft samples were sliced in 2-4 μm thick sections, deparaffinized in xylene and rehydrated through decreasing concentrations (100%, 95%, 90%, 80%, and 70%) of ethyl alcohol, then rinsed in distilled water. Antigen retrieval was performed in 0.01 mol/L Sodium Citrate buffer (pH 6.0) in a microwave oven or in 90°C solution of 0,001 mol/L EDTA for 20 minutes. Endogenous peroxidase activity was quenched with 3% hydrogen peroxide in distilled water. Slides were treated with 1% bovine serum albumin (BSA) (SantaCruz), and 5% normal goat serum in PBS containing 0.05% Triton (Sigma-Aldrich, Inc) and incubated in a closed humid chamber overnight at 4°C with the following antibodies: anti-GLAST (1:200 dilution, Novocastra #NCL-EAAT1); anti-Ki67 (1:50; BD Bioscience #MA5-14520); anti-DCX (1:100; Abcam #ab207175); anti-NEDD9 (1:50, OriGene #TA319569). Staining was detected using the EnVision + System-HRP Labelled Polymer Anti-Rabbit or Anti-Mouse for 1 hour at room temperature and then the Chromogen DAB/substrate reagent (Dako, Glostrup, Denmark). Slides were counterstained with haematoxylin (Sigma Aldrich, Inc), dehydrated and mounted. Quantitative analyses were performed on three to five independent fields per tumor by counting the number of cells in the photographed fields using the 40× objective of a Leica DM-LB microscope. The tumor sections were also stained with hematoxylin and eosin to assess the volume of tumor growth.

Fluorescence studies were performed on GBM specimens provided by the Unit of Pathology at Istituto Besta. For GBM slides immunofluorescence staining was made with: anti-CD31 antibody (1:50, Dako, Glostrup Denmark #GA610), anti-Na+/K+-ATPase α2 (1:50, Millipore #07-674), anti-GLAST (1:100; Novocastra #NCL-EAAT1 or Abcam #ab416), anti-xCT (1:250; Abcam #ab37185). The Alexa Fluor-488 goat anti-rabbit (1:100 dilution, ThermoFisher Scientific #A-1108) and the Cy3-conjugated affine pure goat anti-mouse (1:100 dilution, Jackson ImmuneResearch, Inc.#115-165-003) were used to detect the primary antibodies; cell nuclei were counterstained with DAPI (Sigma Aldrich, Inc.#D9542) and the slides were mounted with a PBS/Glycerol solution. All antibody validation information is available in the product’s manual.

### Magnetic Resonance Imaging / Spectroscopy

MRI/MRS studies were performed on a 7 Tesla BioSpec 70/30 USR scanner equipped with a 12-cm inner diameter actively shielded gradient system reaching a maximum amplitude of 400 mT/m. (Bruker BioSpin, Ettlingen, Germany). The spectra were analyzed with LCModel software (Version 6.3) (Provencher, 1993) for glutamate absolute quantification.

Mice were anaesthetized with 1.5–2% isoflurane (60:40 N_2_O:O_2_ (vol:vol), flow rate 0.8 L/min). To detect the depth of anaesthesia and the animal health condition during the MRI study, the respiratory rate was monitored by a pneumatic sensor. Mice were positioned on an animal bed equipped with a nose cone for gas anaesthesia and a three point-fixation system (tooth-bar and ear-plugs). A 75 mm birdcage linear coil was used for radio frequency excitation and a mouse brain surface coil was used for signal reception. For anatomical references, T2-wi were acquired in three orthogonal planes: axial, sagittal, and coronal. Mice injected with GLAST-enriched or depleted GL261 weekly underwent MRI investigation with the following protocol: high resolution axial T_2_-weighted images (T_2_-wi) acquired using a rapid acquisition with relaxation enhancement (RARE) sequence (TR/TE=3000/13.5 ms, matrix size=256×256, RARE factor=8, slice thickness=0.8 mm, FOV=2.2×2.2 cm^2^, in plane resolution=86 μm, number of averages (NA)=8, acquisition time (AT)=9 min 30 s); 1H spectra carried out by a PRESS sequence (Point RESolved Spectroscopy, TR/TE = 5000/13 ms, adjustment of the first and second order shims conducted beforehand by Mapshim macro of Paravision 5.1 software) from a single voxel (∼10µL) located inside a tumor (or, before detecting solid tumor, at the injection site) and symmetrically in the contralateral hemisphere; contrast agent induced signal enhancement, interpreted as being due to a BBB lesion, was highlighted by 2D T_1_-weighted (T_1_-w) sequence (TR/TE=390/11 ms, matrix sixe=256×256, slice thickness=1 mm, FOV=3×3 cm^2^, in plane resolution=117 μm, NA=4, AT=6 min 36 s); acquired before and after intraperitoneal administration of contrast medium (Gadolinium DTPA).

### Statistical analysis

Cumulative survival curves were constructed using the Kaplan-Meier method and a log-rank test was performed to assess statistical significance between groups (GraphPad PRISM 5.03). Statistical comparisons of the data sets were performed using a two-tailed Student’s T-test, and the results were considered significant when P < 0.05. The experimental groups were defined as reported in (Fleiss *et al*, 2003). No animals or data points were excluded from the analyses for any reason. Blinding and randomization were performed in all experiments.

## Acknowledgments

Thank to Dr. Samanta Mazzetti for helpful hints and suggestion with the confocal microscope. This project was supported by grants from Associazione Italiana per la Ricerca sul Cancro (AIRC) to SP (IG-2013 n. 14323), Fellowship AIRC id. 16557 to CC, a grant from Fondazione Giovanni Celeghin to GF, Fondazione IRCCS Istituto Neurologico C. Besta 5X1000 funds, Il Fondo di Gio and Associazione Italiana Tumori Cerebrali (AITC).

## Author contributions

Designing research studies: SP, GF; Acquiring and analyzing data: SP, CC, NDI, IZ, MC, FP, MP; Writing original draft: SP; Writing review and editing: GF, MGB; Conducting experiments: SP, CC, NDI, IZ,; Supervision, SP, GF, MGB, BP, PLP.

The authors declare that they have no conflict of interest

The paper explained

## Problem

GBM is the most aggressive primary brain tumor: with present treatments overall survival is 15 months. Brain edema and invasive capacity are major determinants of GBM morbidity and lethality. High levels of glutamate are associated to both features and are crucial in favoring GBM infiltration and diffusion by invading normal brain tissue long distance.

The Na+-dependent glutamate transporters including GLAST are expressed by astrocytes and play a relevant role in glutamate uptake and are responsible for maintaining the intracellular levels of glutamate and preventing glutamate-mediated excitotoxicity in the CNS. Their activity depends on the Na+ electrochemical gradient generated by Na+/K+-ATPase. In GBM the glutamate metabolism and trafficking are profoundly altered.

## Results

We studied a large number of GBM specimens and found relevant GLAST expression in 50% of patients in correlation with poor survival. By analyzing GBM stem-like cells growing in vitro as neurospheres and derived-GBM in vivo, we observed a novel and relevant role for GLAST, based on an altered uptake of glutamate associated with a down-regulation of Na+/K+-ATPase. We used for the first time on glioma cells the selective GLAST inhibitor UCPH-101 and showed that it induces apoptosis in GBM stem-like cells but not in astrocytes. A single intratumoral injection of UCPH-101 increased survival in glioma-bearing mice without signs of toxicity. Finally, we described that STAT3 activation is mechanistically related with GLAST up-regulation, suggesting that glutamate can be the master regulator of GLAST expression through STAT3 activation.

## Impact

Increased understanding of this process and the identification of key regulators of GLAST expression and function constitute a preliminary background for translational research leading to the identification of novel GLAST inhibitors to be tested in phase I studies.

